# Metformin Shows Greater Potential Than Semaglutide in Reducing Alzheimer’s Risk in Diabetes Type II via Dual Actions: Tackling Disease Pathways and Environmental Herpesvirus Triggers

**DOI:** 10.1101/2025.03.14.643306

**Authors:** Andrea Georgiou, Panos Zanos, Anna Onisiforou

## Abstract

Alzheimer’s disease (AD) and Diabetes Mellitus Type II (DM2) share overlapping pathological mechanisms, with DM2 increasing AD risk. Disease-modifying therapies (DMTs) for DM2, including Metformin and Semaglutide, have been explored for neuroprotection, yet their efficacy in AD remains unclear. Here, we introduce a novel integrative framework combining comparative network pharmacology, Mendelian Randomization (MR), and transcriptomic validation to systematically assess the therapeutic impact of 39 DM2 DMTs in preventing AD in DM2. Metformin emerged as the top-ranked therapy, whereas Semaglutide ranked among the least effective based on comparative analysis within the DM2-AD pathway-pathway comorbidity network. Notably, a two-sample MR analysis finds no evidence supporting a genetic liability to either drug being associated with AD risk, suggesting that their benefits may arise through non-direct mechanisms or that their relationship could be confounded by third factors. Metformin’s neuroprotective impact is mediated through AMPK, insulin, and adipocytokine signaling, which regulate key AD-related processes. Additionally, Metformin may indirectly affect herpesviruses, emerging environmental contributors to AD, potentially enhancing its neuroprotective effects. In contrast, Semaglutide, despite its growing clinical prominence as a weight loss therapy, exhibits minimal engagement with core neurodegenerative pathways within the DM2-AD comorbidity network, highlighting variability in neuroprotective potential across DM2 DMTs. Furthermore, specific dual-action therapies (e.g., Insulin Glargine and Lixisenatide, Insulin Degludec and Liraglutide) exhibit efficacy comparable to Metformin, reinforcing the need for a precision medicine approach. These findings challenge the assumption that all DM2 DMTs confer equal neuroprotection, revealing significant differences in their impact on AD-related pathways. While some show strong potential for AD prevention, others appear far less effective. Metformin’s effects depend on genotype, disease state, and environmental factors, underscoring the need to re-evaluate DM2 DMTs for AD prevention in population-specific clinical trials. Metformin stands out as a strong candidate for targeted investigations in DM2 patients at high risk of AD.

## 1. Introduction

Alzheimer’s Disease (AD) is a progressive neurodegenerative disorder that currently lacks effective pharmacotherapies ^1^. Affecting over 55 million people globally, AD is the most prevalent form of dementia in the elderly population^2^. AD has been termed “type 3 diabetes” by many researchers due to shared pathological mechanisms with Diabetes Mellitus Type II (DM2) ^3–6^, a metabolic disorder resulting from pancreatic islet β cell failure and insufficient insulin levels for normal glucose metabolism ^7,8^. Individuals with DM2 exhibit a higher incidence of cognitive decline, predisposing them to various forms of dementia, including AD ^9^. Despite the well-documented connection between DM2 and AD, the pathophysiological mechanisms linking metabolic dysfunction in DM2 to increased susceptibility to AD remain poorly understood.

Disease-modifying therapies (DMTs) for DM2 have been proposed as potential pharmacotherapies for AD^10^. DM2 DMTs, such as Metformin and Semaglutide, are currently undergoing clinical trials to evaluate their potential roles in preventing cognitive decline associated with AD ^11,12^. However, while these therapies show promise, the precise molecular mechanisms by which they exert their protective effects remains unclear ^13^. Experimental and preliminary clinical evidence, including observational-studies and randomized clinical trials, suggests that Metformin, a widely used DM2 therapy, may reduce the risk of AD and comorbid AD in individuals with DM2 while also enhancing cognitive performance ^14–19^. Additionally, genetic evidence indicates that Metformin may promote healthy aging ^20^. For instance, an observational study from the National Alzheimer’s Coordinating Center database found that Metformin use was associated with better memory performance compared to non-users, although this improvement was not observed in individuals carrying the APOE-ε4 allele^21^.

Mechanistic studies further reveal Metformin’s impact on AD pathology, influencing factors such as neuronal loss, neural dysfunction, Aβ depositions, tau phosphorylation, and chronic neuroinflammation ^16^. However, existing evidence remains inconsistent and, at times, contradictory ^16^. For instance, in the PDAPP (J9) mouse model of AD, Metformin treatment improved memory function in female mice but worsened it in males, highlighting the importance of sex-specific considerations in evaluating its cognitive effect ^22^. These findings emphasize the need for sex-stratified clinical trials to better understand how Metformin influences cognitive function in men and women. Similarly, population-specific effects have been reported, as a systematic review analysis found that among Asians, Metformin users had a significantly higher risk of AD compared to non-users ^23^. This suggests that genetic, metabolic, or environmental factors may differentially modulate Metformin’s neuroprotective or neurotoxic effects across populations. Such findings reinforce the need for precision medicine approaches when considering Metformin as a therapeutic candidate for AD prevention. Beyond Metformin, other DM2 DMTs, such as GLP-1 analogues, Sodium-Glucose Co-Transporter-2 (SGLT-2) inhibitors, Insulin, Glinides, and Thiazolidinediones, have also shown potential neuroprotective effects in AD ^10,11^. These therapies may have prophylactic effects, particularly in individuals with pre-existing DM2, a known risk factor for AD ^24^. However, their comparative efficacy in reducing AD risk remains unclear, as do the mechanisms through which different types of DM2 DMTs exert neuroprotection.

A major challenge in selecting DMTs for DM2 is that patients respond differently to treatment due to genetic variability, lifestyle influences, and the inherent heterogeneity of DM2. Moreover, confounding factors, including genetic predisposition and aging, further complicate treatment strategies by increasing susceptibility to AD. These variations in treatment response significantly impact the ability of DM2 DMTs to mitigate AD risk, underscoring the importance of identifying key pathways through which these therapies modulate both DM2 and AD pathogenesis.

One critical gap in current research is the influence of environmental factors, such as chronic viral infections, on the efficacy of DM2 DMTs in mitigating AD risk. Viral infections, particularly members of the *Herpesviridae* family, have being implicated in AD onset and progression ^25–28^. Interestingly, Metformin has demonstrated antiviral properties, including the ability to reduce replication of herpesviruses ^29,30^. Recent studies further associate Metformin use with a significantly reduced risk of herpes zoster and postherpetic neuralgia in DM2 patients, with higher cumulative doses providing greater protection ^31^. These results suggest that Metformin’s benefits extend beyond glycemic control, potentially reducing infection-related complications in DM2 patients and mitigating AD risk by addressing viral triggers. However, the interaction between Metformin’s antiviral properties and these environmental factors remains poorly understood. Notably, most ongoing clinical trials investigating Metformin’s role in AD prevention fail to account for environmental contributors, such as chronic viral infections, despite substantial emerging evidence linking them to AD pathology ^25,32,33^. This gap in research highlights the need for integrative approaches that incorporate both genetic and environmental risk factors to fully elucidate Metformin’s neuroprotective effects.

Computational methodologies, such as network pharmacogenomics, pathway enrichment analyses, and the reconstruction of pathway-pathway comorbidity networks, have provided valuable insights into the pathological mechanisms underlying diseases, including comorbidities and the influence of environmental factors such as viral and bacterial infections in disease development and/or progression ^34–39^. Additionally, transcriptomic data analysis can validate predicted drug-pathway interactions by identifying disease-associated gene expression changes in relevant patient populations ^35,40^. By integrating transcriptomic findings with network-based predictions, these approaches refine our understanding of DM2 DMTs in preventing comorbid AD in DM2, offering a more comprehensive framework for assessing their therapeutic potential. These tools enable a systematic investigation of how Metformin modulates pathways linked to metabolic dysfunction, chronic viral infections, and AD, uncovering mechanistic insights that remain unexplored in experimental and clinical studies. This integrative approach paves the way for more targeted and effective interventions.

One significant limitation in the study of comorbid conditions is the lack of transcriptomic data from human patients and animal models of DM2-AD comorbidity in public repositories such as ArrayExpress ^41^ and Gene Expression Omnibus (GEO) ^42^. To address this challenge, we previously developed a novel pathway network-based approach that identifies shared molecular pathways and potential therapeutic targets for comorbidities through the reconstruction and analysis of Disease-Disease pathway-pathway comorbidity networks ^34^. Building on this model, we integrate network pharmacology and graph theory to rank the therapeutic impact of DM2 DMTs, including Metformin, on AD-related pathways within a DM2-AD comorbidity network (see **Figure 1**). To further assess their efficacy, we conducted a Mendelian Randomization (MR) analysis to determine whether Metformin and Semaglutide, the highest- and lowest-ranking DM2 DMTs within the DM2-AD pathway comorbidity network identified through our network pharmacology approach, influence AD risk via genetic mechanisms. We also investigate the therapeutic potential of Metformin combination therapies and its antiviral effects on herpesvirus infections, an emerging environmental factor linked to AD. To validate our findings, we employ two complementary approaches: (1) Blood transcriptomic analysis of samples from patients with mild cognitive impairment (MCI) to compare predicted drug-pathway interactions with real-world gene expression changes. (2) First-neighbor pathway analysis to assess Metformin’s functional connectivity within AD-related processes within the DM2-AD comorbidity network, reinforcing its potential role in AD prevention. This study introduces a novel integrative framework that combines network pharmacology, MR, and transcriptomic validation, providing a data-driven approach to identify DM2 DMTs with the highest potential for AD prevention in DM2 and bridging the gap between computational predictions and clinical relevance. By incorporating network-based drug ranking with genetic and transcriptomic evidence, our approach provides a more comprehensive assessment of therapeutic potential than traditional analyses. Our findings demonstrate that not all DM2 therapies offer equal neuroprotective benefits, highlighting the need for precision medicine strategies that integrate metabolic, genetic, and environmental factors to optimize treatment selection for individuals at risk of comorbid AD in DM2.

**Figure 1:**
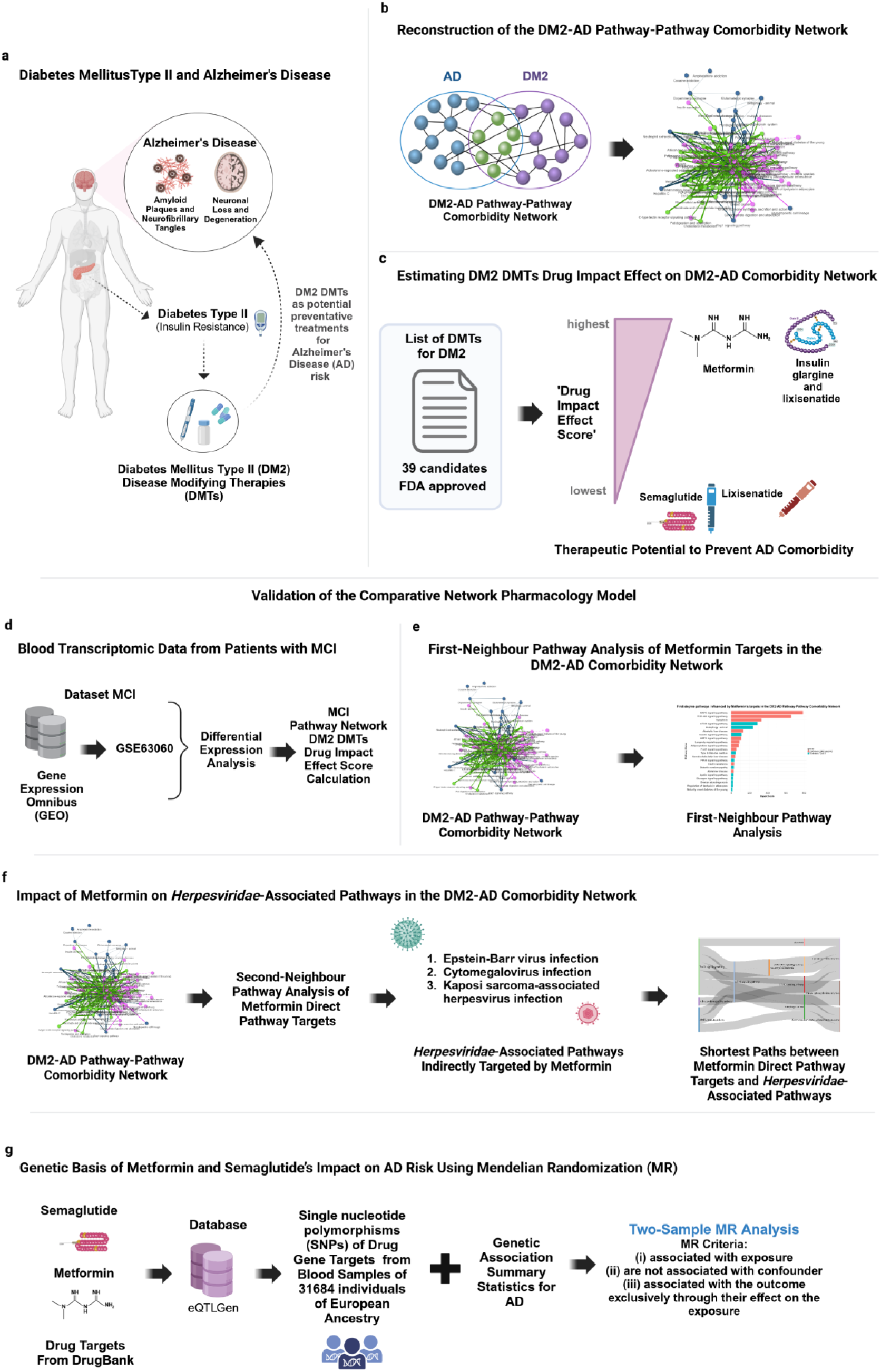
**a.** DM2 increases dementia risk, including AD. Recent findings suggest that certain DM2 DMTs may reduce comorbid AD risk, however existing evidence regarding the effectiveness of the different DM2 DMTs in preventing AD risk is currently unclear and, at times, contradictory. Here, we developed a novel pathway network-based pharmacology approach to perform comparative effectiveness analysis. **b.** We first reconstructed the weighted DM2-AD pathway-pathway network and calculated the ’Disease Node Impact Score’ for each disease-related pathway, which allowed to assess the significance of each node in facilitating the comorbidity between DM2 and AD. **c.** We estimated the Drug Impact Effect Score of DM2 DMTs on AD comorbidity risk by integrating network pharmacology and centrality-based metrics within the DM2-AD pathway comorbidity network. This analysis allowed for the ranking of DM2 DMTs based on their predicted neuroprotective potential, identifying therapies with the highest impact on AD-related pathways. **d.** We validated our network pharmacology model using blood transcriptomic data from patients with MCI, a preclinical stage of AD. Differential expression analysis was performed on the publicly available transcriptomic dataset GSE63060 to identify gene expression changes in MCI patients. The impact of DM2 DMTs was then evaluated by integrating these findings with the MCI-associated pathway network, allowing us to assess how drug-pathway interactions align with real-world molecular alterations. **e.** First-neighbor pathway analysis was conducted within the DM2-AD comorbidity network to examine Metformin’s direct interactions with AD-related pathways. This analysis identified key molecular connections, reinforcing Metformin’s relevance in modulating neurodegenerative processes and its potential role in AD prevention. **f.** To further explore Metformin’s role in AD prevention, we analyzed its indirect influence on *Herpesviridae*-associated pathways, given the emerging link between viral infections and AD risk. A second-neighbor pathway analysis was performed to identify connections between Metformin’s direct targets and pathways involved in Epstein-Barr virus infection, cytomegalovirus infection, and Kaposi sarcoma-associated herpesvirus infection. This revealed potential antiviral mechanisms that may contribute to Metformin’s neuroprotective effects. **g.** A two-sample MR analysis was performed to evaluate whether genetic proxies for Metformin and Semaglutide are associated with AD risk. SNPs from eQTLGen and DrugBank were used as genetic instruments for drug target expression, while genetic summary statistics for AD were obtained from a European ancestry cohort. MR criteria ensured that selected SNPs were associated with drug target expression but not with confounders. This analysis aimed to determine whether Metformin and Semaglutide influence AD risk via genetic mechanisms. ***Figure*** *created with BioRender.com*.

## 2. Results

### 2.1 Network Pharmacology Analysis Highlights Metformin and Dual Therapies as Leading Candidates for Preventing DM2-Associated AD

By developing and applying a novel network pharmacology model, we conducted a comparative effectiveness analysis of DM2 DMTs to assess their potential suitability in preventing comorbid AD in DM2. Unlike traditional approaches that evaluate single-disease mechanisms, our network-based framework integrates multi-pathway interactions, capturing shared and distinct molecular processes underlying DM2-AD comorbidity. This addresses key gaps in existing comorbidity studies, which often overlook complex disease interactions and fail to systematically rank therapeutic candidates based on their functional impact across interconnected pathways. This was achieved by constructing and analyzing a weighted DM2-AD pathway-pathway comorbidity network, consisting of 129 nodes and 536 edges (**see Figure 1b**). The network includes 70 shared pathways between AD and DM2, 28 AD-specific pathways, and 31 DM2-specific pathways (Supplementary Tables S1-S2).

To evaluate the role of each pathway in facilitating comorbidity, we calculated the ’Disease Node Impact Score’ for each disease-related pathway. This enabled us to quantify its significance in the DM2-AD comorbidity network. A crucial aspect of our analysis was determining the therapeutic potential of DM2 DMTs, which we achieved by calculating the ’Drug Impact Effect Score’ for each therapy. This score was derived by summing the ’Disease Node Impact Score’ of all pathways targeted by a given DMT within the DM2-AD comorbidity network. Finally, we ranked DM2 DMTs based on their ’Drug Impact Effect Score’, identifying those with the highest therapeutic potential for reducing AD comorbidity risk in individuals with DM2 (**see Figure 1c**).

Using the DrugBank ^43^ and the Kyoto Encyclopedia of Genes and Genomes (KEGG) ^44^ databases, we collected 57 DM2 DMTs. Drug-pathway associations were available for 50 of these therapies, while seven DMTs (Pramlintide acetate, Dapagliflozin, Canagliflozin, Empagliflozin, Ertugliflozin, Luseogliflozin hydrate, and Colesevelam hydrochloride) lacked pathway data in KEGG and were excluded from pathway-based calculations. For the 50 DMTs with available data, we identified 85 drug-pathway associations, which were used to calculate the ’Drug Impact Effect Score’ within the DM2-AD pathway-pathway comorbidity network. Drugs targeting pathways outside the comorbidity network were assessed only for their relevant interactions, while those exclusively targeting pathways outside the network were excluded. Ultimately, from the 50 DMTs, only 39 were found to interact with pathways within the DM2-AD comorbidity network and were included in the final ranking.

Our analysis identified Metformin as the highest-impact DM2 DMT in preventing or reducing comorbid AD in DM2, outperforming 38 other therapies interacting with pathways within the DM2-AD comorbidity network (see **Figure 2**). Metformin’s therapeutic benefits appear to be mediated through modulation of AMPK, Insulin, and Adipocytokine signaling pathways. Additionally, combination therapies such as Insulin Glargine & Lixisenatide and Insulin Degludec & Liraglutide demonstrated impact scores comparable to Metformin, suggesting that these dual-action therapies may provide enhanced protection against comorbid AD in DM2. While insulin analogs (Insulin Glargine and Insulin Detemir) ranked among the top 10 DM2 DMTs, their impact scores remained lower than these specific combination therapies, highlighting the potential advantage of multi-target approaches.

**Figure 2:**
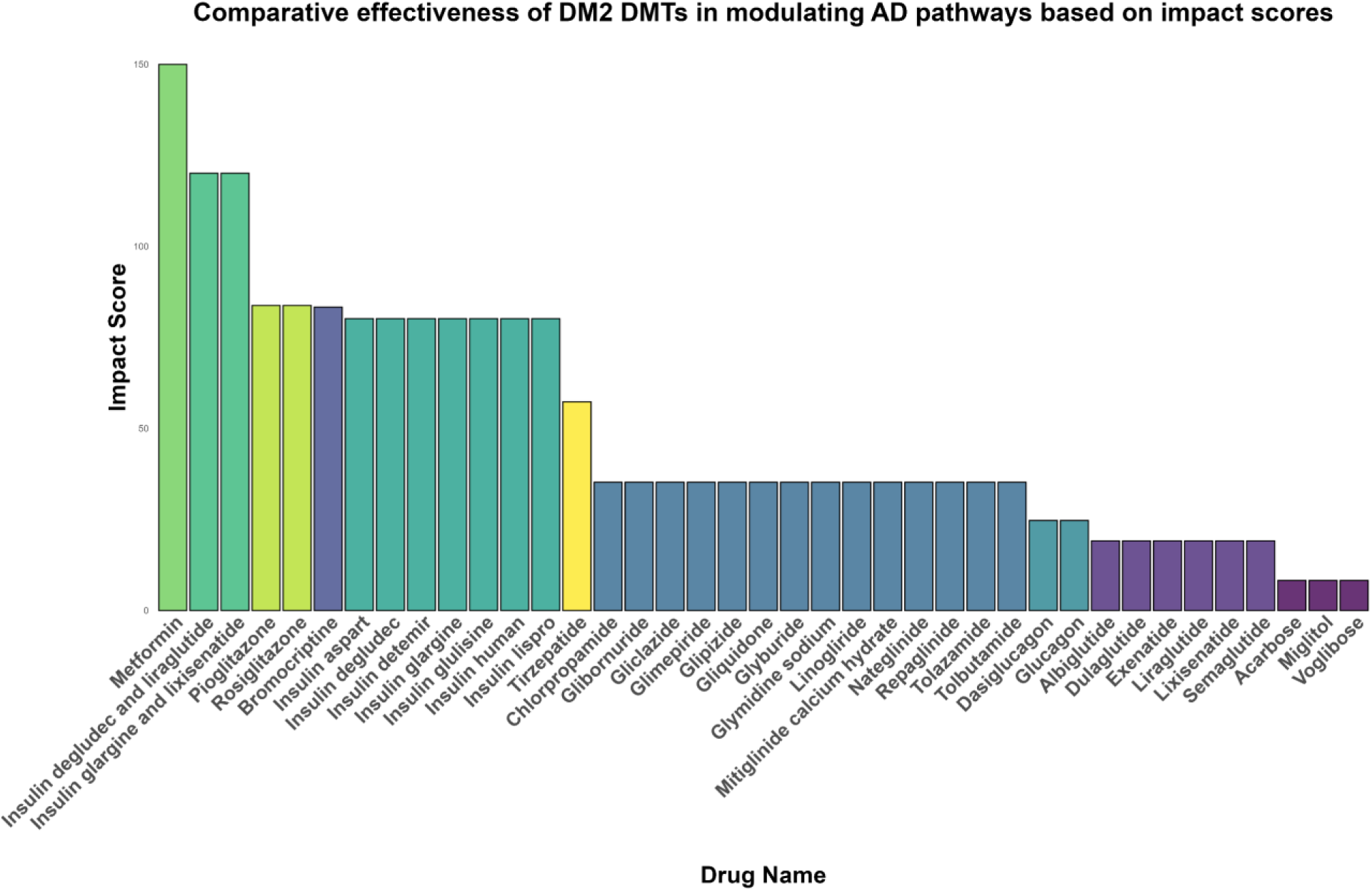
Comparative effectiveness of DM2 DMTs in the DM2-AD comorbidity network based on impact scores. Drugs with identical scores are represented by the same color.

Furthermore, Pioglitazone and Rosiglitazone ranked among the top 10 DM2 DMTs with high impact scores for AD prevention, exerting their effects through modulation of the PPAR and AMPK signaling pathways. Our computational model also identified Bromocriptine as a high-impact DM2 DMT, influencing AD-related pathways through the Neuroactive ligand-receptor interaction, Dopaminergic synapse, Prolactin signaling, and Parkinson’s Disease pathways. Conversely, Semaglutide, along with Lixisenatide, Liraglutide, Exenatide, Dulaglutide, and Albiglutide, ranked among the lowest-impact DM2 DMTs, primarily targeting the Neuroactive ligand-receptor interaction and Insulin secretion pathways. The lowest impact scores were observed for Acarbose, Miglitol, and Voglibose, which mainly influence the Carbohydrate digestion and absorption pathway.

### 2.2 Metformin Combination Therapies with Higher Impact in Preventing Comorbid AD in DM2

We also investigated Metformin combination therapies with the highest impact scores in preventing comorbid AD in DM2, evaluating their effects on pathways within the DM2-AD comorbidity network. Our analysis identifies several Metformin combinations with high impact scores, suggesting enhanced potential for reducing AD risk in DM2 patients.

Among these, Metformin combined with Rosiglitazone maleate (KEGG ID: D10244) and Metformin combined with Pioglitazone hydrochloride (KEGG ID: D09744) stand out, both exhibiting an impact score of 233.754, the highest among Metformin combination therapies. Thiazolidinediones (Rosiglitazone/Pioglitazone) are Peroxisome Proliferator-Activated Receptor (PPAR)-γ agonists, known for their role in improving insulin sensitivity and modulating metabolic and inflammatory pathways. This drug combination modulates multiple pathways within the DM2-AD comorbidity network, including AMPK signaling (hsa04152), PPAR signaling (hsa03320), Insulin signaling (hsa04910), and Adipocytokine signaling (hsa04920). Since both Metformin and thiazolidinediones independently regulate AMPK signaling, this combination reinforces AMPK’s key role in metabolic regulation and neuroprotection.

Other Metformin-based combinations, including Glipizide & Metformin (KEGG ID: D10265), Glyburide & Metformin (KEGG ID: D10266), and Metformin & Repaglinide (KEGG ID: D10500), exhibit a moderate yet substantial impact score of 185.2094. Glipizide, Glyburide, and Repaglinide are sulfonylurea receptor agonists that enhance insulin secretion by stimulating pancreatic β-cells. The combination of Metformin and sulfonylureas modulates multiple pathways within the DM2-AD comorbidity network, including Insulin secretion (hsa04911), AMPK signaling (hsa04152), Insulin signaling (hsa04910), Adipocytokine signaling (hsa04920), and the Type II diabetes mellitus pathway (hsa04930). This combination likely contributes to AD prevention via improved insulin sensitization and glucose regulation, albeit with a lower impact than thiazolidinedione-based combinations, which additionally engage PPAR signaling alongside AMPK regulation.

Additionally, Glipizide, Glyburide, and Repaglinide also target the ABC transporters pathway (hsa02010), which is involved in cellular transport mechanisms and drug metabolism. Although not included in the DM2-AD comorbidity network, this interaction with sulfonylureas may indicate off-target effects related to drug transport and clearance.

### 2.3 Validation of Model-Predicted DM2 DMT Effects on AD Prevention in DM2

To assess the reliability of our network pharmacology model predictions, we applied two independent validation strategies. The first approach utilized blood transcriptomic data from patients with MCI to evaluate whether the predicted drug impact effects aligned with real-world gene expression changes. The second approach employed network-based validation through first-neighbor pathway analysis, examining whether Metformin’s direct pathway targets were functionally connected to AD-associated pathways within the DM2-AD comorbidity network.

#### 2.3.1 Validation of DM2 DMTs in AD Prevention Using Blood Transcriptomic Data from Patients with MCI

To validate our comparative network pharmacology model, we analyzed blood transcriptomic data from patients with MCI, an early stage of AD. Since DM2 DMTs primarily act peripherally, particularly in the pancreas, and MCI precedes AD, analyzing blood transcriptomic changes in MCI patients provides insights into early molecular alterations associated with AD development and progression. Furthermore, identifying DM2 DMT-associated transcriptomic signatures in blood may uncover systemic mechanisms through which these therapies contribute to AD prevention.

To validate our model predictions, we mapped DM2 DMT-targeted pathways onto enriched pathways derived from differentially expressed genes (DEGs) in MCI patients, assessing whether the predicted drug impact scores from the comparative analysis of DM2 DMTs within the DM2-AD comorbidity network aligned with real-world transcriptomic data. This validation allowed us to test the robustness of our model and the predictive value of our ranking approach. For this analysis, we utilized blood transcriptomic data from MCI patients (GSE63060 dataset), comparing them to healthy controls to identify DEGs and enriched pathways. We then constructed a KEGG pathway-pathway network based on these findings and mapped DM2 DMT-pathway interactions to determine which DM2 therapies had the highest impact scores in MCI. By integrating our model predictions with an independent transcriptomic dataset, we provided an additional validation layer for our ranking system, reinforcing its potential utility in identifying DM2 DMTs with therapeutic relevance in AD prevention.

Differential expression analysis identified 6,628 DEGs (adjusted p-value < 0.05). Enrichment analysis revealed 37 statistically significant KEGG pathways (Supplementary Table S3), including pathways targeted by DM2 DMTs, such as AMPK signaling (hsa04152), Protein digestion and absorption (hsa04974), Neuroactive ligand-receptor interaction (hsa04080), and Insulin resistance (hsa04931). Pathway comparison between the MCI and DM2-AD networks revealed 18 overlapping pathways, suggesting a shared molecular basis for neurodegeneration in both conditions (**Figure 3**). However, 111 pathways were unique to the DM2-AD network, indicating additional processes driving AD in DM2 patients. Conversely, 19 pathways were exclusive to MCI, potentially reflecting neurodegenerative processes independent of metabolic dysfunction.

**Figure 3:**
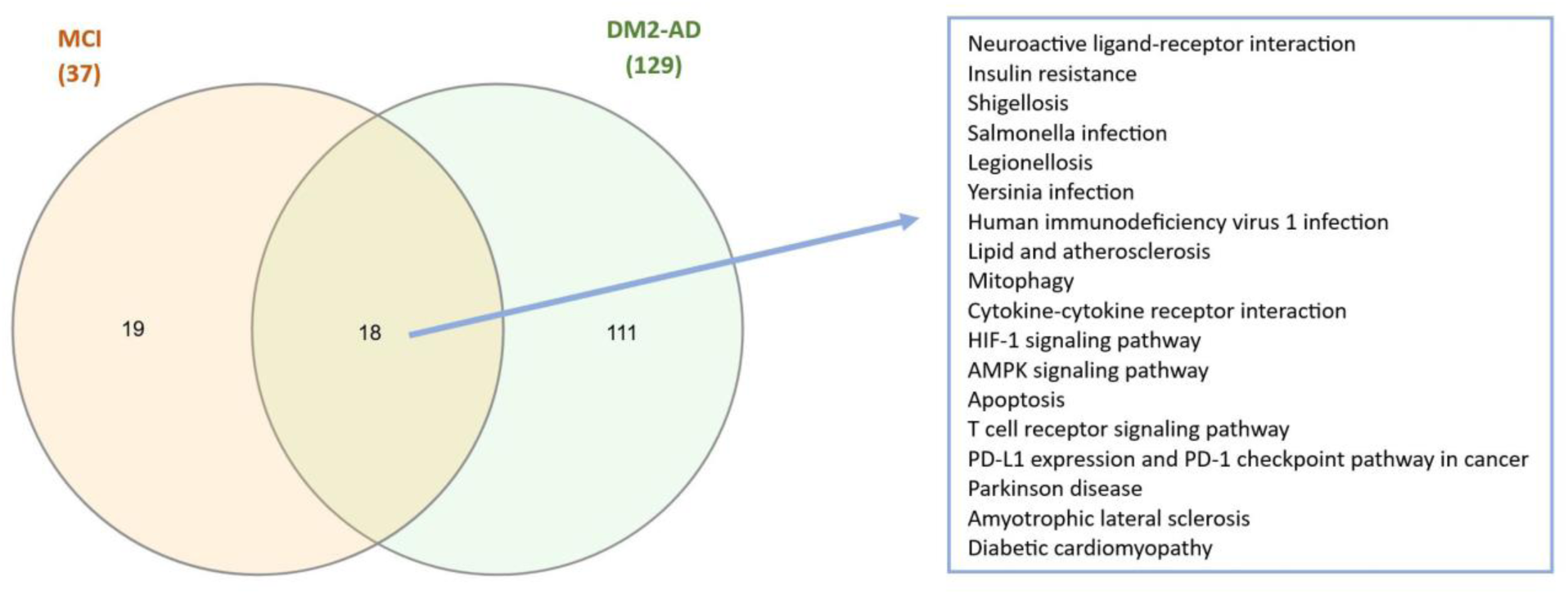
Venn diagram comparing pathways between the MCI network and the DM2-AD pathway network.

The comparative effectiveness analysis demonstrated that the highest-ranking drugs in the DM2-AD comorbidity pathway network (**Figure 2**) also ranked among the highest in the MCI pathway network (**Figure 4**), reinforcing the robustness of our ranking approach. More specifically, Bromocriptine, Metformin, Pioglitazone, and Rosiglitazone, identified among the top five drugs in the DM2-AD pathway comorbidity network, were also among the top five in the MCI pathway network, with minor variations in ranking position. In the MCI network, Bromocriptine ranked first and Metformin second, whereas in the DM2-AD comorbidity network, Metformin ranked first and Bromocriptine fifth. These variations suggest that while both models consistently identify key DM2 DMTs with high impact scores, differences in ranking arise due to differences between pathways involved in MCI and DM2-induced AD. Drugs primarily targeting metabolic dysfunction, such as Metformin, ranked higher in the DM2-AD network, while neuroactive agents, such as Bromocriptine, ranked higher in the MCI network. This suggests that the effectiveness of DM2 DMTs in AD prevention may depend on the underlying disease driver, whether metabolic or related to neuronal dysfunction. Nonetheless, the overlap in top-ranking drugs across both networks strengthens confidence in their potential role in AD prevention, as these drugs consistently target pathways central to neurodegeneration and metabolic dysfunction.

**Figure 4:**
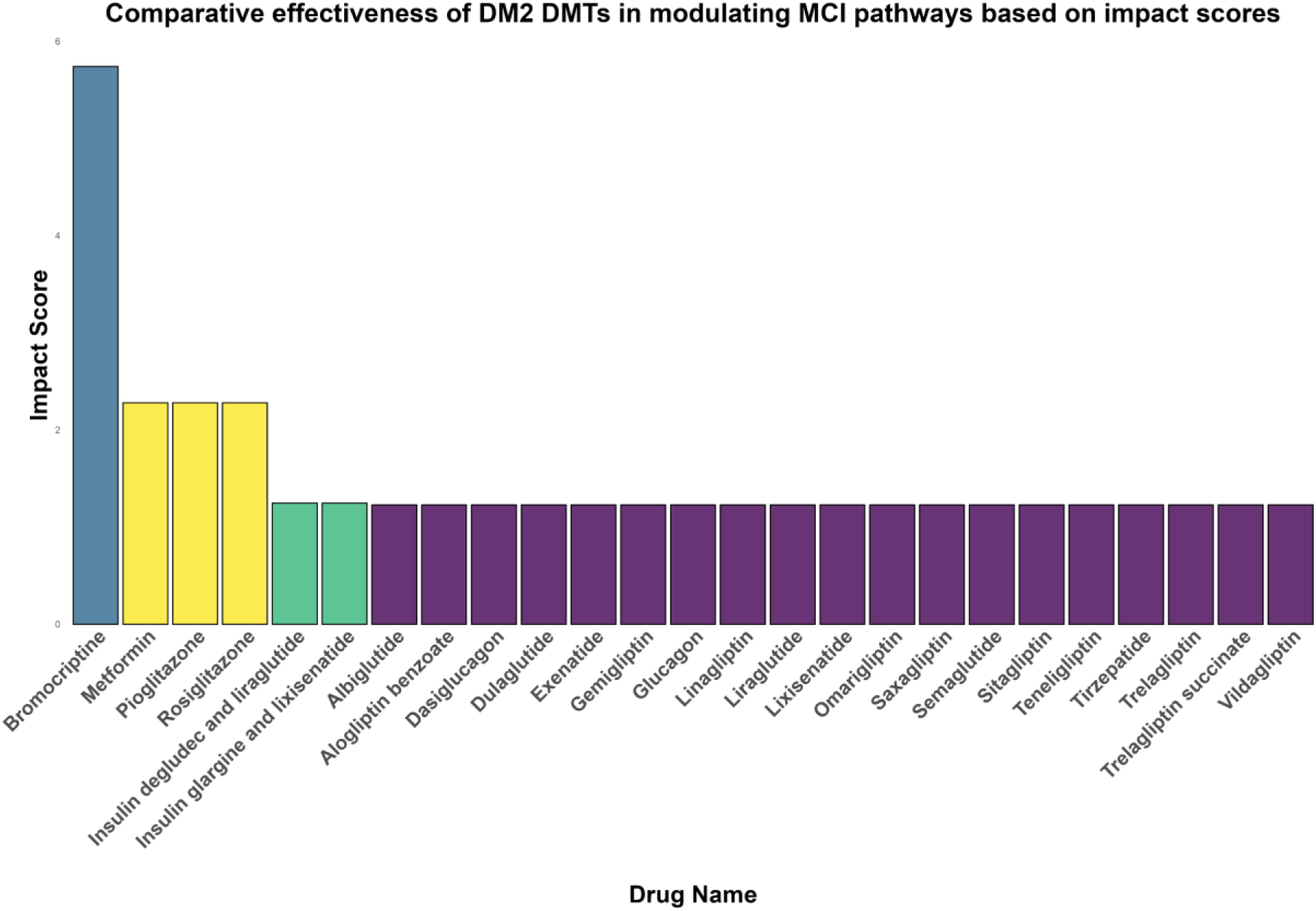
Comparative effectiveness of DM2 DMTs in modulating pathways within the MCI network based on impact scores. Drugs with identical scores are represented by the same color.

These results emphasize the importance of evaluating drug impact across different conditions to determine which therapies have the strongest potential for AD prevention in specific patient populations. The overlap in top-ranked drugs across both networks further supports the predictive value of our network pharmacology model in identifying DM2 DMTs with neuroprotective potential in DM2.

#### 2.3.2 Network-Based Validation of Metformin’s Effects within the DM2-AD Comorbidity Network Using First-Neighbor Pathway Analysis

To further validate Metformin’s predicted effects in the DM2-AD comorbidity network, we examined whether its direct targets were functionally linked to AD-associated pathways. First-neighbor pathway analysis revealed that Metformin’s primary targets, including the AMPK signaling pathway (hsa04152), Insulin signaling pathway (hsa04910), and Adipocytokine signaling pathway (hsa04920), are directly connected to pathways involved in AD pathogenesis, reinforcing its potential neuroprotective effects.

Notably, the Alzheimer’s disease pathway (hsa05010) was identified as a first-degree neighbor of the Insulin signaling pathway (hsa04910) targeted by Metformin, highlighting a direct molecular link between Metformin’s metabolic effects and neurodegenerative mechanisms (**Figure 5**). Additionally, several first-neighbor pathways associated with both AD and DM2, including the Apoptosis pathway (hsa04210), Longevity regulating pathway (hsa04211), and PI3K-Akt signaling pathway (hsa04151), were identified. The Longevity regulating pathway (hsa04211) is influenced through Metformin’s effects on the Insulin signaling pathway (hsa04910) and Adipocytokine signaling pathway (hsa04920). These pathways are critical for cell survival, inflammation, and metabolic regulation, further supporting Metformin’s role in AD prevention.

**Figure 5:**
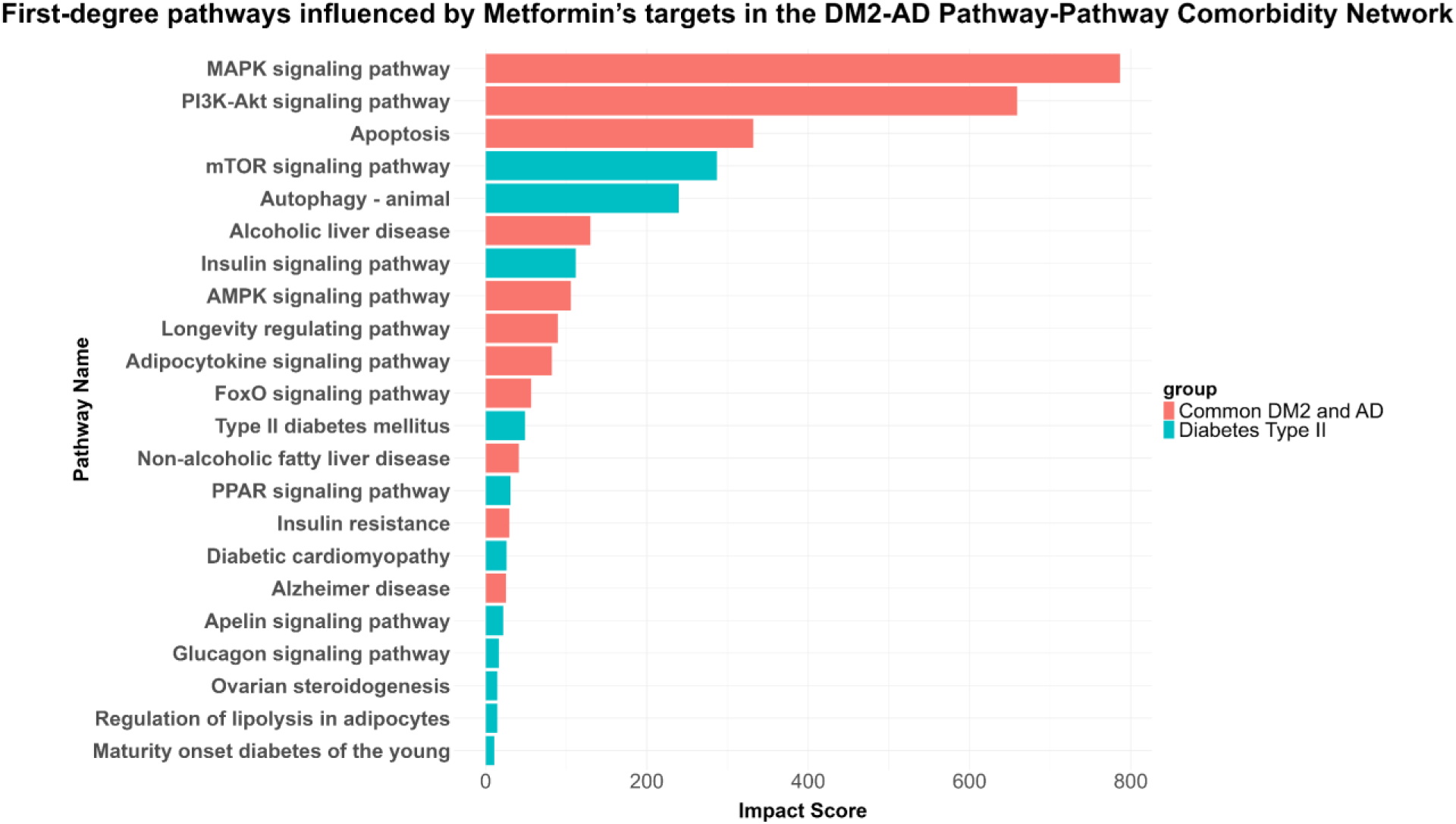
First-degree pathways influenced by Metformin’s targets in the DM2-AD Pathway Comorbidity Network.

The presence of the longevity pathway suggests that Metformin’s effects extend beyond glucose metabolism, potentially influencing aging and neuroprotection. This analysis provided an additional validation layer, confirming that Metformin-targeted pathways interact with AD-related pathways within the DM2-AD comorbidity network. The connectivity of Metformin’s targets bridges metabolic dysfunction in DM2 with neurodegenerative processes in AD, reinforcing its systemic impact on disease-modifying mechanisms. By integrating first-neighbor pathway analysis, we strengthened confidence in Metformin’s ranking within our network pharmacology model and its potential for repurposing in AD prevention.

### 2.4 Metformin’s Effects on Herpesviridae-Associated Pathways in the DM2-AD Comorbidity Network

We investigated whether Metformin’s effects in the DM2-AD comorbidity network extend to pathways associated with *Herpesviridae* infections, given the growing evidence linking viral infections, particularly herpesviruses, to an increased risk of AD ^25,28,32,33^. Notably, several *Herpesviridae*-associated pathways, including Epstein-Barr virus infection (hsa05169), Cytomegalovirus infection (hsa05163), and Kaposi sarcoma-associated herpesvirus infection (hsa05167), were identified as part of the DM2-AD comorbidity network. These pathways are shared between DM2 and AD, suggesting a potential role of viral mechanisms in both neurodegeneration and metabolic dysfunction.

To explore Metformin’s indirect influence on these pathways, we analyzed second-degree interactions of its pathway targets within the DM2-AD comorbidity network. Specifically, we identified second-degree neighbors of Metformin’s primary targets, including the AMPK (hsa04152), Insulin (hsa04910), and Adipocytokine (hsa04920) signaling pathways. Our analysis revealed that Metformin’s targets can indirectly influence the Cytomegalovirus (hsa05163) and Kaposi sarcoma-associated herpesvirus (hsa05167) pathways through their network interactions (**Figure 6**). Additionally, the Adipocytokine and Insulin signaling pathways can indirectly modulate the Epstein-Barr virus infection pathway (hsa05169), suggesting a potential link between Metformin and viral mechanisms associated with AD.

**Figure 6:**
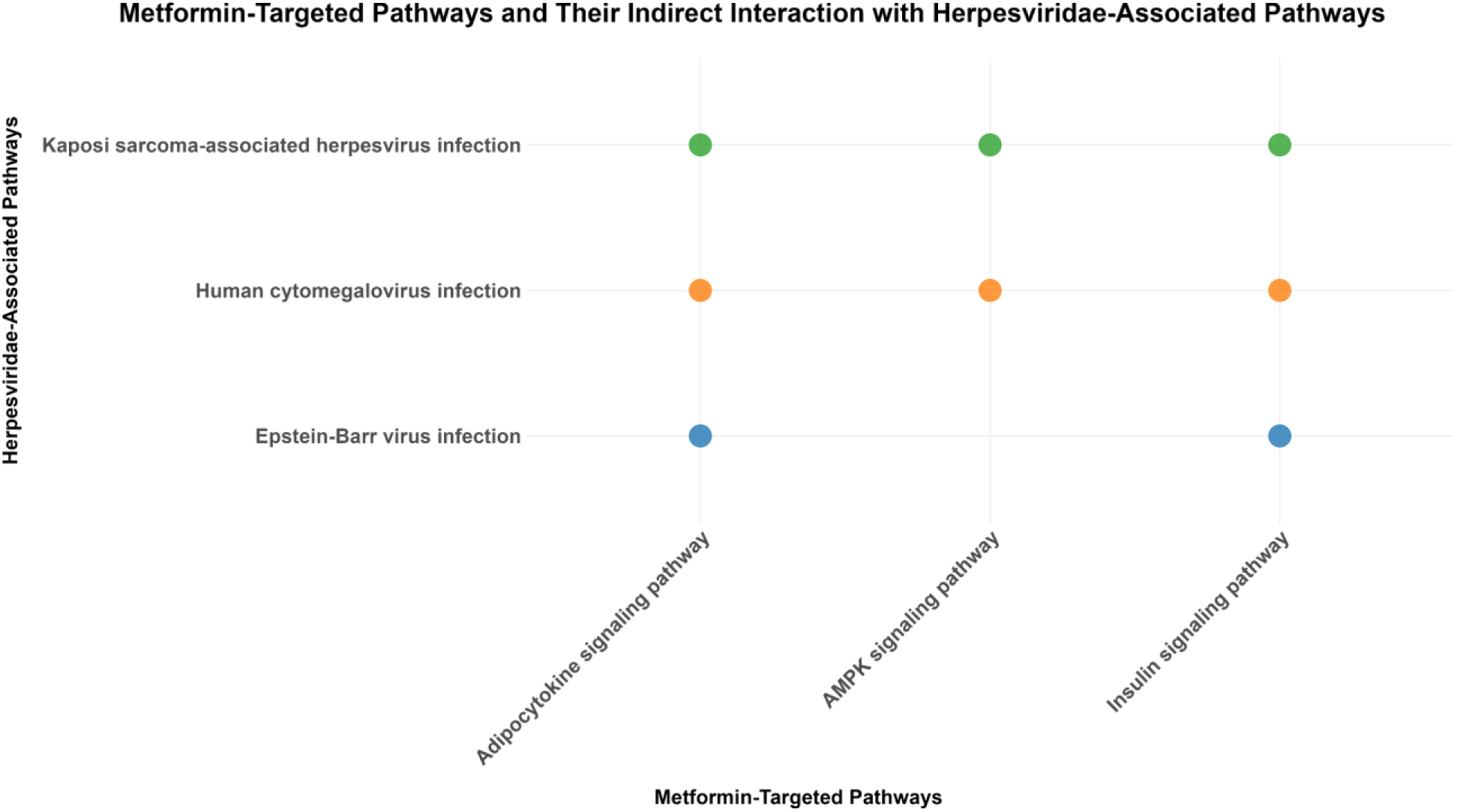
Indirect Influence of Metformin-Targeted Pathways on Herpesviridae-Associated Pathways in the DM2-AD Comorbidity Network.

These findings suggest that Metformin’s beneficial effects in DM2-AD comorbidity may extend beyond metabolic regulation and neuroprotection, potentially modulating viral pathways implicated in AD pathogenesis. To further investigate the mechanistic links between Metformin and *Herpesviridae*-associated pathways, we extracted all shortest paths connecting Metformin-targeted pathways to these viral pathways within the DM2-AD comorbidity network (**Figure 7**). The shortest path analysis identified key molecular bridges through which Metformin’s pathway targets may indirectly influence viral pathways associated with Epstein-Barr virus, Cytomegalovirus, and Kaposi sarcoma-associated herpesvirus infections.

**Figure 7:**
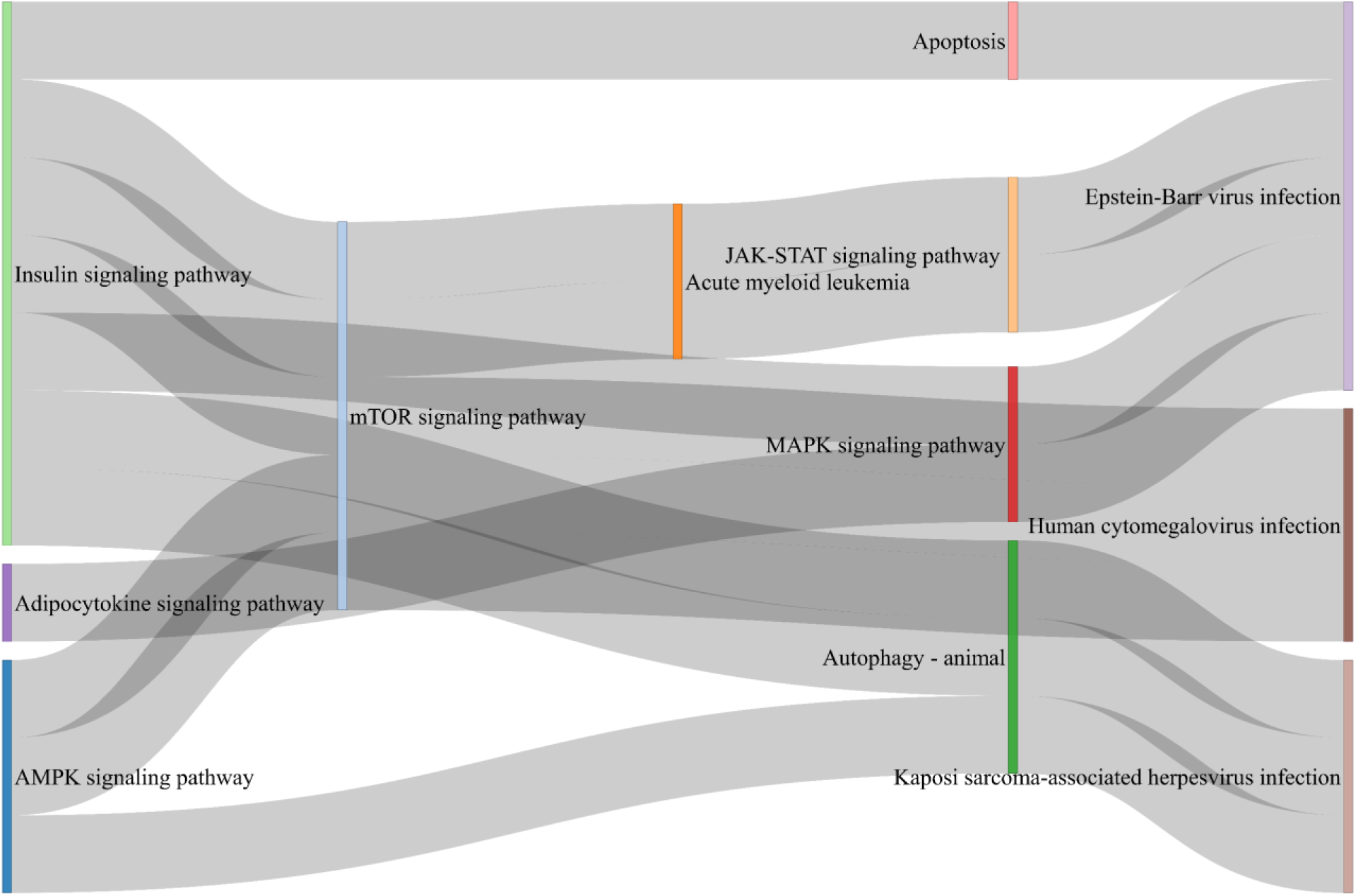
Sankey Diagram illustrating the shortest paths linking Metformin-primary targeted pathways to Herpesviridae-associated pathways within the DM2-AD Comorbidity Network.

Several notable patterns emerged from this analysis. First, the AMPK signaling pathway was linked to Epstein-Barr virus infection via the mTOR and JAK-STAT signaling pathways, both of which play critical roles in immune regulation and cellular stress responses. This suggests that Metformin’s activation of AMPK may have downstream effects on viral infection pathways, potentially modulating inflammatory and metabolic responses linked to neurodegeneration. Similarly, the Insulin signaling pathway was connected to Epstein-Barr virus infection through the MAPK and Apoptosis pathways, as well as to Cytomegalovirus infection via the mTOR pathway. Given that both insulin resistance and viral infections have been implicated in AD pathology, these interactions further support the hypothesis that Metformin may exert neuroprotective effects by influencing immune-metabolic cross-talk.

The Adipocytokine signaling pathway, another key Metformin target, was found to interact with Epstein-Barr virus infection through the MAPK pathway and with Cytomegalovirus and Kaposi sarcoma-associated herpesvirus infections via the Insulin signaling and Autophagy pathways. The presence of autophagy-related interactions is particularly noteworthy, as autophagy dysfunction has been implicated in both viral persistence and AD pathogenesis. This suggests that Metformin’s ability to enhance autophagy could contribute to both viral suppression and neuroprotection. By identifying the shortest molecular routes linking Metformin’s targets to viral pathways implicated in AD, we highlight a potential mechanism through which Metformin may mitigate AD risk via modulation of viral infections.

### 2.5 Investigating the Genetic Basis of Metformin and Semaglutide’s Impact on AD Risk Using MR

To complement our pathway-based analysis, we conducted a MR analysis to examine whether the genetic proxies for Metformin and Semaglutide’s target gene expression levels are causally associated with AD risk. These drugs were selected due to their ongoing clinical trials for AD, with Metformin emerging as the highest-ranking drug in our pathway-based analysis and Semaglutide ranking among the five lowest. The MR approach aimed to test the causal relationship between their genetically proxied effects and AD risk, providing a genetic perspective to complement the mechanistic insights identified in our pathway analysis.

We conducted a two-sample MR analysis that indicated no overlap between the exposure (Metformin and Semaglutide drugs) and outcome (AD) samples. The MR analysis did not support a causal association between the two genetically proxied drugs effects and AD risk (Metformin-AD: IVW OR = 1.00, 95% confidence interval (CI): 0.96 to 1.05, P=0.94; Semaglutide-AD: IVW OR = 1.02, 95% CI: 0.99 to 1.04, P=0.11) (**Table 1**). Additionally, sensitivity analysis found no evidence of pleiotropy (MR–Egger intercept p > 0.05; MR–PRESSO Global Test p > 0.05). However, high heterogeneity was observed in the Semaglutide-AD association (Cochran Q =55.85, P < 0.05), suggesting variability in the genetic instruments, which may affect the reliability of the results for this drug (**Table 1**). All retained SNPs demonstrated F-statistics greater than 10, confirming their strength as instrumental variables for the genetically proxied drugs, Metformin and Semaglutide (Supplementary Tables S4-S6).

**Table 1:** Results of inverse variance-weighted MR and sensitivity analysis for Metformin/Semaglutide drugs and risk of AD.

| Exposure | Outcome | No SNPs | IVW: OR (95%CI) | MR-Egger: OR (95%CI) | MR-PRESSO p-value | Cochran's Q p-value |
| --- | --- | --- | --- | --- | --- | --- |
| Metformin | AD | 28 | 1.02 (0.99-1.04) | 1.00 (0.97-1.04) | 0.12 | 0.06 |
| Semaglutide | AD | 29 | 1.00 (0.96 - 1.05) | 1.02 (0.94-1.11) | 0.16 | 0.001 |

Overall, the MR analysis did not provide evidence that the genetic proxies for Metformin or Semaglutide influence AD risk. This does not rule out their potential protective effects but indicates that such effects, if present, may not operate through the genetic variants analyzed in this study. The high heterogeneity observed for Semaglutide further underscores the challenges of using genetic proxies for this drug. In contrast, the pathway-based analysis identified Metformin as the highest-impact DM2 DMT in preventing or reducing comorbid AD in DM2, based on its interactions within the DM2-AD comorbidity network. Additionally, transcriptomic validation demonstrated that Metformin’s predicted pathway interactions align with real-world gene expression changes in MCI patients, reinforcing the biological plausibility of its neuroprotective effects. These findings suggest that Metformin’s influence on AD risk may be mediated through non-genetic mechanisms rather than through direct genetic determinants. Together, these findings demonstrate that while genetic proxies do not support a direct causal effect of Metformin or Semaglutide on AD risk, pathway-based and transcriptomic analyses provide complementary evidence suggesting a biologically relevant role for Metformin in AD prevention.

## 3 Discussion

The current study provides a comparative evaluation of DM2 DMTs in preventing AD comorbidity in DM2. By integrating network pharmacology, MR, and transcriptomic validation, we systematically assessed the therapeutic impact of these treatments. Our computational model ranks DM2 DMTs within the DM2-AD comorbidity network based on their Drug Impact Effect Score, offering valuable guidance for clinicians and researchers in prioritizing therapies that may lower AD risk in DM2 patients. However, pathway-based scores serve as a proxy rather than definitive proof of clinical efficacy. While they provide a mechanistic framework for identifying high-potential drugs based on their interactions within disease-associated networks, they do not replace direct clinical validation. To address this, we incorporated MR and transcriptomic validation, which provide complementary evidence beyond network pharmacology. Although MR analysis did not establish a genetic causal link between Metformin and reduced AD risk, transcriptomic validation confirmed that Metformin’s predicted pathway interactions align with real-world gene expression changes in MCI patients, reinforcing its biological relevance. Ultimately, while pathway-based scoring identifies promising candidates, clinical efficacy must be confirmed through longitudinal studies and randomized clinical trials. Our study presents a data-driven strategy for prioritizing therapies, guiding future experimental and clinical research in AD prevention for DM2 patients.

Our findings suggest that Metformin has a greater predicted neuroprotective impact compared to 38 other DM2 DMTs, including Semaglutide, that target the DM2-AD comorbidity network, primarily by engaging key metabolic and neuroinflammatory pathways and demonstrating antiviral properties against herpesviruses. Evidence supports Metformin’s antiviral properties, with studies linking its use to reduced herpesvirus replication and a lower risk of herpes zoster and postherpetic neuralgia in DM2 patients, particularly at higher cumulative doses ^29–31^. In contrast, Semaglutide, despite its widespread clinical use for glycemic control and weight loss, showed minimal interaction with neurodegenerative pathways within the DM2-AD comorbidity network, suggesting that its metabolic benefits may not extend to AD risk reduction. While Semaglutide is highly effective for weight loss, Metformin has modest weight-reducing effects, primarily through its role in stimulating the production of N-lactoyl-phenylalanine, an appetite-suppressing metabolite generated in the intestine ^45^. Individuals with DM2 taking Metformin typically experience a ≥5% reduction in initial body weight within the first year ^46^, whereas Semaglutide treatment achieves 15% or more weight loss, as demonstrated in the OASIS 1 Clinical Trial ^47^.

GLP-1 receptor agonists like Semaglutide regulate appetite and glycemic control primarily by activating the GLP-1 receptor (GLP-1R), which is part of the Neuroactive Ligand-Receptor Interaction Pathway. This pathway comprises neuropeptides, neurotransmitters, and hormone receptors that mediate intercellular communication, particularly in the central nervous system (CNS) and endocrine signaling. However, in our analysis, the Neuroactive Ligand-Receptor Interaction Pathway did not exhibit direct connectivity with the rest of the pathways within the DM2-AD comorbidity network, suggesting that GLP-1 receptor activation may have a limited direct role in neurodegenerative processes.

Thus, while GLP-1 receptor agonists like Semaglutide are widely recognized for their metabolic benefits, their role in neuroprotection remains uncertain. Although some GLP-1RAs can cross the blood-brain barrier (BBB) and may exert neuroprotective effects ^48^ , the extent and precise mechanisms of their impact on neurodegenerative processes remain unclear. Thus, it is important to distinguish whether their neuroprotective potential arises from direct CNS actions or from peripheral metabolic improvements that indirectly benefit the brain. Preclinical studies suggest that Semaglutide may have therapeutic potential in neurodegenerative disorders like AD and Parkinson’s disease (PD) ^49^. In AD models, Semaglutide reduces amyloid-beta plaques, mitigates neuroinflammation, and improves cognitive and behavioral deficits ^49^. In PD models, it enhances mitochondrial function, promotes neurogenesis, and protects dopaminergic neurons, improving motor function. These findings highlight Semaglutide’s potential disease-modifying effects ^49^ Despite preclinical evidence supporting Semaglutide’s neuroprotective effects, our pathway analysis reveals that the Neuroactive Ligand-Receptor Interaction Pathway does not directly integrate into the DM2-AD comorbidity network. Thus, while Semaglutide may exert some degree of neuroprotection, its effects are not sufficient to significantly mitigate AD pathology in the context of DM2. Given these findings, further research is needed to determine whether Semaglutide’s effects on AD are primarily mediated through metabolic improvements or if there are yet unidentified direct CNS mechanisms contributing to neuroprotection.

Additionally, weight loss alone does not inherently equate to cognitive resilience, particularly when it results from appetite suppression rather than adaptive metabolic reprogramming. Unlike caloric restriction, which triggers adaptive metabolic reprogramming, including enhanced autophagy and neuronal plasticity ^50,51^, Semaglutide primarily reduces food intake through appetite suppression ^52^. While this promotes weight loss, it may not activate the same neuroprotective pathways. Instead, it carries risks of nutritional deficiencies^53^ and muscle loss^54^, both of which are risk factors for cognitive decline. These findings underscore the importance of distinguishing between metabolic therapies that actively modulate neuroprotective pathways, such as Metformin, and those that primarily function through systemic appetite suppression. Future studies should investigate whether GLP-1 receptor agonists confer long-term cognitive benefits beyond their weight loss effects and whether these effects are sustained across diverse patient populations.

In contrast, Metformin appears more effective in preventing AD in DM2 patients by targeting broader metabolic and neuroprotective pathways, including AMPK, insulin, and adipocytokine signaling, which are directly implicated in AD pathogenesis. AMPK activation decreases apoptosis, inflammation and oxidative stress and inhibits insulin resistance ^55^, all of which are essential for protecting neurons from AD-related degeneration ^56^. These effects help mitigate β-amyloid accumulation, tau hyperphosphorylation, and neuroinflammation, reinforcing Metformin’s potential role in preventing AD comorbidity in DM2 patients. Our first-neighbor pathway analysis further highlights direct molecular connections between Metformin’s targets and AD-associated pathways, including the Alzheimer’s disease pathway, Apoptosis pathway, and Longevity regulating pathway. Additionally, to further validate our comparative network pharmacology model, we analyzed blood transcriptomic data from patients with MCI, an early stage of AD. Our analysis revealed that the highest-ranking drugs in the DM2-AD comorbidity pathway network also ranked among the top drugs in the MCI pathway network, with Metformin being the second highest-ranked drug. This alignment supports the robustness of our model in identifying therapeutic candidates with neuroprotective potential and reinforces Metformin’s promise as a therapeutic candidate for AD prevention in DM2 patients.

An emerging area of AD research highlights the role of chronic viral infections, particularly herpesviruses, in neurodegeneration ^25,28,32,33^. Our findings suggest that Metformin’s impact on herpesvirus-associated pathways may contribute to its protective effects in DM2-AD comorbidity. Notably, previous studies have identified significant enrichment of the Epstein-Barr virus infection pathway in female AD patients, particularly within the CA1 hippocampal region, indicating a sex-specific susceptibility to Epstein-Barr virus-driven neurodegeneration ^40^. Furthermore, Metformin has been shown to inhibit the replication of multiple viruses, including Influenza A, and herpesviruses such as Cytomegalovirus and Epstein-Barr virus^30,57^, reinforcing its potential antiviral and neuroprotective benefits. Given the growing evidence linking viral infections to AD, Metformin’s dual action in targeting both metabolic dysfunction and viral-associated pathways suggests a novel therapeutic avenue for mitigating AD risk in DM2 patients. These findings warrant further investigation into Metformin’s antiviral mechanisms in AD pathogenesis, particularly in populations at higher risk for viral reactivation and neurodegeneration.

Our findings indicate that Metformin-thiazolidinedione combinations (Metformin + Rosiglitazone/Pioglitazone) have the highest impact in preventing AD, primarily through AMPK, PPAR, insulin, and adipocytokine signaling pathways. In contrast, Metformin-sulfonylurea combinations (Metformin + Glipizide/Glyburide/Repaglinide) show moderate impact, mainly influencing insulin secretion and glucose metabolism, suggesting that thiazolidinedione-based therapies may offer superior AD protection. Sulfonylureas target ABC transporters (hsa02010), yet these were not identified in the enrichment analysis for AD or DM2, despite their role in drug metabolism, BBB permeability, and β-amyloid clearance^58,59^ . This may reflect a limitation of enrichment analysis, which prioritizes statistically significant pathways while overlooking indirectly associated mechanisms. Future research should examine how sulfonylurea-induced ABC transporter modulation affects drug bioavailability, BBB function, and β-amyloid clearance, potentially influencing AD risk in DM2 patients.

Moreover, our results demonstrate that specific dual action therapies (e.g., Insulin Glargine and Lixisenatide, Insulin Degludec and Liraglutide) exhibit efficacy comparable to Metformin. Notably, evidence suggests that six months therapy with a fixed combination of Liraglutide and Degludec in a group of very old DM2 subjects who responded to treatment showed changes in n Gram-negative *Alistipes* gut microbiota , correlating with cognitive improvement ^60^. Given that gut microbiota imbalances may contribute to AD progression by promoting amyloid-beta accumulation, triggering inflammation, increasing oxidative damage, and impairing insulin signaling ^28,61^, these findings highlight a potential modulation of gut-brain axis may be a key mechanism underlying the cognitive benefits of dual-action therapies in DM2-AD comorbidity. Further research is needed to investigate how these therapies influence microbiota composition and their subsequent impact on brain function.

The results from the MR analysis and pathway-based computational model provide complementary insights into the potential role of Metformin in reducing AD risk. While pathway analysis identified Metformin and certain combination therapies as strong candidates for preventing DM2-associated AD by modulating key pathways, MR analysis found no evidence supporting a causal protective relationship between genetically proxied Metformin effects and AD risk. This divergence suggests that the protective effects identified in the pathway analysis may involve mechanisms beyond genetic proxies, such as metabolic conditions (e.g., insulin resistance, inflammation), cellular states (e.g., energy stress, mitochondrial function), and environmental factors (e.g., antiviral effects, microbiota composition changes). A strength of this study is the use of genetic data from exceptionally large AD GWAS meta-analysis. Another strength is the use of MR analysis, a method able to overcome common sources of bias in observational studies such as residual confounding and reverse causality. However, there was significant heterogeneity between the genetic instruments used as proxies for Semaglutide drug, which suggests that at least some of the variants have pleiotropic effects. However, the MR-Egger intercept showed no evidence of directional pleiotropy and the MR estimates from additional sensitivity analyses were consistent with the null finding, meaning that considerable bias due to pleiotropy is unlikely.

Observational evidence further supports a context-dependent effect of Metformin, as data from the National Alzheimer’s Coordinating Center database found that Metformin use was associated with improved memory performance, but this benefit was not observed in APOE-ε4 carriers ^21^, suggesting that Metformin may be protective in individuals with DM2 but not in those with certain genetic predisposition factors for AD, such as APOE-ε4, which is the strongest genetic risk factor for sporadic AD ^62^. Thus, Metformin’s effects may be insufficient to counteract APOE-ε4-mediated pathology. This highlights the potential need for genotype-specific therapeutic approaches in AD prevention strategies. Additionally, conflicting findings suggest that prolonged Metformin treatment may exacerbate AD pathology in genetically susceptible models, increasing β-amyloid accumulation, tau phosphorylation, and synaptic dysfunction ^63^. Behavioral studies in AD 3xTg-AD transgenic mice have reported memory impairment and enhanced amyloidogenic processing following chronic Metformin administration, raising concerns about its effects in non-DM2 populations ^63^. However, evidence also suggests that Metformin may have cognitive benefits in certain contexts, as it has been shown to enhance attention, inhibitory control, and associative learning in younger non-transgenic C57BL/6 mice (≤16 months) ^63^. These findings highlight the complexity of Metformin’s effects, which may be age-, genotype-, and disease-context dependent, reinforcing the need for further experimental and clinical validation to clarify its therapeutic potential and risks in AD prevention across different populations.

Despite the strengths of our study, several limitations should be acknowledged. First, our analysis does not account for confounding factors such as genetic variability, lifestyle influences, and other comorbidities or concurrent DMTs, which may affect AD risk and treatment response in DM2. Additionally, our network pharmacology approach relies on curated databases that may not capture all relevant biological interactions, and our findings may evolve as knowledge expands. While our MR analysis found no genetic evidence supporting a causal protective effect of Metformin or Semaglutide on AD, this does not rule out non-genetic mechanisms, such as Metformin’s antiviral properties or microbiota modulation, which are not captured by MR analysis. Moreover, as the transcriptomic data of MCI patients are derived from peripheral blood rather than brain tissue, they may not fully reflect CNS-specific molecular changes. However, given that DM2 DMTs primarily modulate peripheral metabolism, their systemic effects, including improved insulin sensitivity and reduced inflammation, may indirectly influence brain health. Another key limitation is the degree to which computational findings translate to clinical applications. While our network-based approach provides valuable insights into potential therapeutic candidates, these predictions require validation through well-designed clinical trials. Additionally, sex-specific effects remain underexplored, as evidence suggests differential susceptibility to Epstein-Barr virus-driven AD in females ^40^, which may interact with DM2 DMTs in influencing AD risk. Future studies should further investigate these factors to refine precision medicine approaches for DM2-associated AD prevention.

Despite these limitations, our findings have important therapeutic implications for clinical decision-making and AD prevention strategies in patients with DM2. Given Metformin’s higher ranking, its potential role in AD prevention should be further explored in clinical trials assessing its long-term cognitive benefits beyond glycemic control. importantly, the context-dependent effects of Metformin, including its reduced benefit in APOE-ε4 carriers and potential risks in non-DM2 populations, emphasize the need for stratified patient selection when considering DM2 DMTs for AD prevention. These findings emphasize the importance of personalized treatment approaches that integrate metabolic, genetic, and environmental factors to optimize therapeutic outcomes. Future clinical guidelines should incorporate precision medicine frameworks that tailor treatment choices based on individual risk factors ^64^, ensuring therapies are optimized for both metabolic health and neuroprotection. Furthermore, our study provides a validated framework for ranking DM2 DMTs based on their predicted impact within the DM2-AD comorbidity network, offering a data-driven approach to identify therapies with the highest potential for AD prevention. By integrating network pharmacology, MR analysis, and transcriptomic validation, we demonstrate that Metformin’s multimodal actions, spanning metabolic regulation, neuroinflammation, and antiviral defense, may underlie its stronger neuroprotective effects. Most importantly, our findings challenge the assumption that all DM2 DMTs provide equal neuroprotection, revealing significant variability in their impact on AD-related pathways. While some therapies show strong potential for AD prevention, others appear far less effective. The effects of DM2 DMTs are influenced by genotype, disease state, and environmental factors, highlighting the need for population-specific clinical trials to re-evaluate their role in AD prevention. Future research should incorporate longitudinal clinical trials, multi-omics integration, and patient stratification to refine therapeutic recommendations for DM2-associated AD prevention.

## 4 Conclusion

This study presents a comparative evaluation of DM2 DMTs in preventing DM2-associated AD comorbidity by integrating network pharmacology, MR analysis, and transcriptomic validation. While the findings highlight differences in how these therapies engage neurodegenerative pathways, they should be interpreted within the scope of computational modeling. The ranking of therapies is based on their pathway-targeting effectiveness within the DM2-AD comorbidity network, and while Metformin demonstrates the highest engagement with key neuroprotective mechanisms, and Semaglutide among the least, this does not imply complete inefficacy of lower-ranked therapies. Instead, the results suggest that some DM2 DMTs act through indirect, metabolic, or context-dependent mechanisms rather than direct neuroprotection. This study does not provide causal clinical evidence but rather a framework for prioritizing therapies for further research. Future clinical validation, particularly in diverse populations considering genetic, metabolic, and environmental factors, is necessary to confirm these computational findings.

## 5 Methods

### 5.1 Comparative Network Pharmacology Effectiveness Analysis of DM2 DMTs in Preventing AD Comorbidity in DM2

#### 5.1.1 Reconstruction of the DM2-AD Pathway-Pathway Comorbidity Network

To assess the therapeutic potential of DM2 DMTs in preventing comorbid AD in DM2, we first reconstructed the DM2-AD pathway-pathway comorbidity network, following an established methodology from previous work ^34^. This approach enables the identification of shared molecular mechanisms and potential therapeutic targets in comorbid diseases through the construction of disease-disease pathway-pathway networks ^34^. By using the *STRING disease* app in Cytoscape, we first collected the top 200 disease-associated proteins with the highest disease scores for DM2 (DOID:9352) and AD (DOID:10652). *STRING: disease* acquires its data from the DISEASES database ^65^, which integrates gene-disease associations from multiple sources, including automatic text mining, manually curated databases, and genome-wide association studies. Confidence scores are assigned to these associations, ranging from 1 star (low confidence) to 5 stars (high confidence). Next, we performed enrichment analysis on the 200 disease-associated proteins of AD and DM2 using the Kyoto Encyclopedia of Genes and Genomes (KEGG) database. ClueGO ^66^ an app in Cytoscape, was used to conduct the enrichment analysis, considering only statistically significant pathways (adjusted p-value ≤ 0.05, Bonferroni step-down correction).

To construct the functional relationships between pathways, we utilized the KEGGREST package ^44^ in R to parse the KEGG database (Homo sapiens, accessed 02/02/2025). This process extracted 1923 functional pathway-pathway interactions from 363 KEGG pathways. We then integrated these interactions with the significantly enriched pathways obtained from DM2 and AD disease-associated proteins, forming the DM2-AD pathway-pathway comorbidity network. The final DM2-AD comorbidity network consists of 129 pathways (nodes) and 536 edges, representing functional interactions between pathways. Among these, 70 pathways are shared between AD and DM2, highlighting common molecular mechanisms.

Additionally, we assigned weights to the edges of the constructed comorbidity network to reflect their biological significance. Specifically, edges involving interactions between intersection nodes, which are pathways that can be found in both diseases, were assigned a weight of 3. Additionally, edges that involve interactions between an intersection and non-intersection nodes received a weight of 2. Edges that represent functional interactions between DM2 and AD pathways were also assigned a weight of 2. All other nodes were assigned a weight of 1. This weight assignment aimed to prioritize interactions between nodes from the two diseases that play a more significant role in facilitating the emergence of their comorbidity.

The weighting scheme was designed to prioritize interactions with higher biological relevance in comorbidity formation. While no universal standard exists for assigning edge weights in comorbidity networks, our approach follows established practices in network medicine, where higher weights are given to interactions between nodes that are functionally or mechanistically critical ^67,68^.

#### 5.1.2 Estimating DM2 DMTs Drug Impact Effect Scores in the DM2-AD Comorbidity Network

To identify which DM2 DMTs may have the greatest therapeutic potential for preventing AD comorbidity risk in DM2, we developed a novel network theory-based methodology to estimate drug impact on a comorbidity network. Drugs were ranked from highest to lowest based on their therapeutic potential for reducing AD risk in DM2. Our approach builds on prior studies in network medicine that have demonstrated the utility of centrality-based metrics for identifying key pathways in disease comorbidity networks. Network-based drug repurposing has leveraged betweenness and closeness centrality to rank pathway significance and predict drug-disease associations ^69,70^ , while centrality metrics have also been used to examine disease co-occurrence and mechanistic overlaps ^34,71^. Expanding on these principles, our study applies network pharmacology to systematically rank DM2 DMTs based on their predicted impact on AD prevention in DM2. This data-driven framework prioritizes therapies within the DM2-AD comorbidity network, offering a novel strategy to identify drugs with the highest potential for reducing AD risk in DM2 patients.

### Calculation of Disease Node Impact Scores

To estimate the impact of each drug, we first calculated the ‘Disease Node Impact Score’ for each pathway (node) within the DM2-AD pathway-pathway network. The Disease Node Impact Score for a vertex v was determined using the following equation:

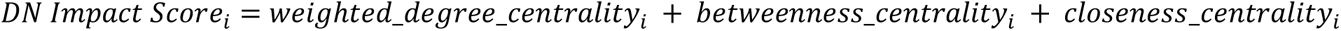

The *weighted_degree_centrality* represents the total weight of edges connected to each vertex in the network. The *betweenness_centrality* quantifies how frequently a vertex acts as a bridge along the shortest paths between other vertices in the graph, taking into account the edge weights and normalizing the results. Lastly, the *closeness_centrality* assesses how close a vertex is to all other vertices in the graph, while also considering the edge weights and normalizing the result.

This combined measure aims to capture the importance or influence of vertices (disease-related pathways) in the comorbidity network. It considers different aspects such as direct connections (weighted degree centrality), bridging roles (betweenness centrality), and overall closeness to other vertices (closeness centrality) to comprehensively assess the significance of each pathway in facilitating the emergence of this comorbidity. Nodes were ranked based on their Disease Node Impact Scores, from highest to lowest.

Nodes within the comorbidity network that were disconnected (i.e., lacking edges or connections to other nodes) were not initially assigned a score, as centrality metrics depend on network connectivity. Disconnected nodes may result from incomplete pathway interaction data, missing intermediary nodes needed to establish full connectivity, or the inherent lack of interacting pathways for a specific pathway within the network. To ensure their potential contribution was considered, they were assigned the minimum Disease Node Impact Score observed in the DM2-AD comorbidity network. This prevented the exclusion of potentially relevant pathways that may have limited connectivity due to incomplete data rather than a true lack of involvement in DM2-AD comorbidity. By incorporating all pathways into the analysis, this approach ensures a more comprehensive assessment while acknowledging limitations in current pathway interaction data.

### Calculation of Drug Impact Effect Scores

We then calculated the Drug Impact Effect Score for DM2 DMTs in the DM2-AD pathway-pathway comorbidity network to assess their potential therapeutic impact on AD comorbidity prevention. First, we retrieved the names of FDA-approved, currently used (i.e., not discontinued) drugs for DM2 from DrugBank ^43^ and the KEGG ^44^ databases, specifically from the Type II diabetes mellitus pathway entry (hsa04930) and disease entry H00409. After removing duplicate entries, a total of 57 DM2 DMTs were collected. To calculate the ’Drug Impact Effect Score’ for each DMT in the DM2-AD comorbidity network, we utilized the KEGGREST package ^72^ in R to parse the KEGG database ^44^ and extract drug-pathway associations. Drug-pathway interaction data were available for 50 out of the 57 DM2 DMTs, while seven DM2 DMTs (Pramlintide acetate, Dapagliflozin, Canagliflozin, Empagliflozin, Ertugliflozin, Luseogliflozin hydrate, and Colesevelam hydrochloride) lacked pathway associations in KEGG and were excluded from further analysis.

For the 50 DM2 DMTs, we identified 85 drug-pathway associations from KEGG database. Ultimately, only 39 of these DMTs were found to target pathways within the DM2-AD comorbidity network and were included in the Drug Impact Effect Score calculations. The ’Drug Impact Effect Score’ of each DMT is calculated by summing up the ‘Disease Node Impact Score’ of all the nodes (pathways) that each DMT targets within the comorbidity network. Drugs that targeted both pathways within and outside the comorbidity network were evaluated based solely on their effects on the comorbidity network pathways. Conversely, drugs that exclusively targeted pathways absent from the comorbidity network were omitted from the analysis. Finally, DM2 DMTs were ranked from highest to lowest based on their Drug Impact Effect Score, with higher scores indicating greater therapeutic potential for preventing AD comorbidity risk in DM2.

#### 5.1.3 Calculation of Drug Impact Effect Scores within the DM2-AD Comorbidity Network for Metformin Combination Therapies

We also investigated which Metformin combination therapies exhibit the highest impact scores in preventing AD in DM2. The impact scores for Metformin combination therapies were determined by evaluating their interactions within the DM2-AD comorbidity network. The total impact score for each combination therapy was obtained by summing the individual pathway impact scores, calculated by the ‘Disease Node Impact Score’ (as described in Section 5.1.2) for the pathways modulated by the respective drugs. Combination therapies were identified from the KEGG database under the Type 2 Diabetes Mellitus entry (H00409). To ensure biological relevance, combination treatments that included drugs without targets within the DM2-AD comorbidity network were excluded from the analysis. For example, combination therapies involving Linagliptin and Alogliptin benzoate, both Dipeptidyl Peptidase-4 (DPP-4) inhibitors, were omitted as they primarily target the Protein Digestion and Absorption pathway, which was not part of the comorbidity network and thus did not contribute to the network-based impact analysis. This approach ensured that only relevant drug combinations were assessed for their potential impact on AD prevention in DM2 patients.

#### 5.1.4 Validation of DM2 DMTs in AD Prevention Using Blood Transcriptomic Data from Patients with MCI

To validate the impact of DM2 DMTs, including Metformin, in preventing AD comorbidity in DM2, we analyzed publicly available blood transcriptomic data from patients with MCI, a well-established preclinical stage of AD. We utilized the GSE63060 dataset, which includes 80 MCI cases and 104 controls (PMID: 26343147). Gene expression data were generated with the Illumina HumanHT-12 V3.0 expression beadchip (Platform GPL6947). Pre-processing and analysis were conducted in the R environment for statistical analysis and visualization ^73^. The dataset was normalized and log2 transformed before further processing. To identify DEGs, we used the Limma R package ^74^, which enables robust differential expression analysis in microarray experiments. Probe-set IDs were mapped to their corresponding gene symbols based on the platform’s annotation files. When multiple probe-set IDs corresponded to the same gene, the average expression value was used. Genes with an adjusted p-value < 0.05 were considered statistically significant and selected for further pathway analysis.

We then performed enrichment analysis using the KEGG database and the ClueGO app in Cytoscape, allowing for the identification of statistically significant pathways associated with MCI. Only pathways with an adjusted p-value ≤ 0.05 (corrected using Bonferroni step-down) were considered significant. Next, we constructed the KEGG MCI pathway-pathway network using the statistically enriched pathways identified from the transcriptomic data, following the methodology outlined in Section 5.1.1. We then mapped DM2 DMTs-pathway interactions onto these pathway network, as described in Section 5.1.2, to determine which DM2 DMTs have the highest impact score and validate Metformin’s role in AD prevention.

The Drug Impact Effect Score was calculated as described in Section 5.1.2; however, unlike the DM2-AD network, which was weighted, the MCI pathway network was unweighted. In the DM2-AD network, edge weights were assigned based on the extent of pathway overlap between the two diseases, reflecting their contribution to comorbidity. In contrast, the MCI network relied on an unweighted approach, as pathways were derived solely from DEGs in MCI patients, without a secondary disease context to establish weighted connections. Consequently, drug impact scores were computed using the same centrality-based metrics.

As in the DM2-AD comorbidity network, we accounted for disconnected nodes within the MCI network by assigning them the minimum impact score rather than excluding them. This approach ensured that all pathways remained part of the analysis, allowing for a more comprehensive assessment of drug impact while preventing the loss of potentially relevant but less-connected pathways. Finally, we aggregated the pathway impact scores for each DM2 DMT, ranking drugs based on their total impact within the MCI network. This allowed us to assess whether the highest-ranked drugs in the DM2-AD network also maintained a strong ranking in the MCI network, providing additional validation for the predictive value of our model.

#### 5.1.5 Network-Based Validation of Metformin’s Impact on AD-Related Pathways Using First-Neighbor Pathway Analysis in the DM2-AD Comorbidity Network

To further validate Metformin’s relevance in DM2-AD comorbidity, we performed a first-neighbor pathway analysis within the DM2-AD comorbidity network. This analysis aimed to determine whether Metformin’s direct targets were functionally linked to AD-associated pathways, providing additional support for its potential neuroprotective effects. First, we identified the first-neighbor pathways of Metformin’s primary targets, including the AMPK signaling pathway (hsa04152), Insulin signaling pathway (hsa04910), and Adipocytokine signaling pathway (hsa04920) within the DM2-AD comorbidity network. To map the immediate molecular connections of these pathways, we extracted their first-degree neighbors using the igraph package in R. For each Metformin-targeted pathway, we identified the pathways that are directly connected to them in the DM2-AD comorbidity network. Additionally, for each first-neighbor pathway of Metformin’s targets, we extracted the pathway impact scores within the network, as calculated in Section 5.1.2.

### 5.2 Investigating the Impact of Metformin in *Herpesviridae*-Associated Pathways in the DM2-AD Comorbidity Network

To assess whether Metformin’s effects in the DM2-AD comorbidity network extend to pathways associated with *Herpesviridae* infections, we analyzed its second-degree pathway interactions. Given the growing evidence linking viral infections, particularly herpesviruses, to an increased risk of AD, we aimed to identify indirect connections between Metformin’s primary targets and *Herpesviridae*-associated pathways.

Following a similar approach to Section 5.1.5, we utilized the igraph package in R to identify the second-degree neighbor pathways of Metformin’s primary targets, including the AMPK signaling pathway (hsa04152), Insulin signaling pathway (hsa04910), and Adipocytokine signaling pathway (hsa04920) within the DM2-AD comorbidity network. We then isolated interactions specifically related to Epstein-Barr virus infection (hsa05169), Cytomegalovirus infection (hsa05163), and Kaposi sarcoma-associated herpesvirus infection (hsa05167). These three Herpesviridae pathways were present in the DM2-AD comorbidity network, allowing us to explore potential indirect mechanisms through which Metformin may influence viral-associated pathways linked to AD risk.

To further investigate the mechanistic links between Metformin and Herpesviridae-associated pathways, we extracted all shortest paths connecting Metformin-targeted pathways to these viral pathways within the DM2-AD comorbidity network. Using the all_shortest_paths() function in the igraph package, we computed the most direct network-based routes between Metformin’s primary targets and each of the three *Herpesviridae*-associated pathways.This analysis allowed us to identify all intermediary pathways that may mediate Metformin’s effects on viral processes implicated in neurodegeneration, providing insight into potential indirect mechanisms through which Metformin may influence AD risk.

### 5.3 Mendelian Randomization Analysis of Genetically Proxied Drug Effects on AD

We conducted a two-sample MR study to investigate the causal relationship between two genetically proxied drugs (Metformin and Semaglutide) and AD. We identified the corresponding gene targets for metformin and Semaglutide through DrugBank and selected the genetic variants associated with their corresponding mRNA expression levels (Supplementary Table S4).

Cis-eQTL data for the drug gene targets were downloaded from the eQTLGen database based on the blood samples of 31,684 individuals from European ancestry (www.eqtlgen.org) Single nucleotide polymorphisms (SNPs) located in significant cis-eQTL loci (p < 1 × 10− ^5^) were utilized as the instrumental variables (IVs) for the two drugs. Beta and standard error for each variant were calculated from z-score and p-value, which were provided by the eQTLGen database. To retain the independent IVs, SNPs were clumped and discarded at linkage disequilibrium r2 < 0.001 within a 10,000 kb window, which was based on European ancestry reference data from the 1000 Genomes Project ^75^.

Genetic association summary statistics for AD were retrieved from the largest publicly available GWAS meta-analysis study by Bellenguez *et al*. ^76^. The substantial participant numbers contributed to the enhanced statistical power of the study (111,326 clinically diagnosed/‘proxy’ AD cases and 677,663 controls).

The inverse variance weighted (IVW) method was used as the main MR analysis and the weighted median (WM) and MR-Egger methods as sensitivity analyses. The three methods were based on three different assumptions: (1) the selected SNPs are associated with the exposure; (2) the selected SNPs are not associated with confounders; and (3) the selected SNPs are associated with the outcome exclusively through their effect on the exposure ^77^. The F-statistic was calculated to evaluate genetic instrument strength. MR-PRESSO and Cochran’s Q statistics were used to evaluate pleiotropy and heterogeneity, respectively ^78^.

MR was employed to examine causal associations between drug targets and AD, using the TwoSampleMR package in R ^78^.

## Data Availability

The data used in this study are obtained from publicly available repositories, as referenced in the Methods section.

## Code Availability

The analyses of the study utilized a number of publicly available software packages which can be found through in-text citations. The in-house R script utilized for the comparative effectiveness analysis of the DM2 DMTs on the DM2-AD comorbidity network is available upon request to the corresponding author.

## Ethics declarations

### Ethics approval and consent to participate

Not applicable.

## Acknowledgements

Research was supported by Research & Innovation Foundation of Cyprus – Excellence Hubs 2021 (EXCELLENCE/0421/0543) grant to P.Z., with A.O. being listed as a co-I, and a European Commission Marie Skłodowska-Curie fellowship #101031962 to P.Z. This publication was made possible by support from the IDSA Foundation to P.Z. and A.O. Its contents are solely the responsibility of the authors and do not necessarily represent the official views of the IDSA Foundation.

## Author Contributions

AO conceived and designed the study. AO and AG performed the analysis and drafted the original manuscript. AO, AG, and PZ contributed to manuscript review, editing, and refinement. All authors have read and approved the final version for publication.

## Supplementary Information

